# SARS-CoV-2 cellular coinfection is limited by superinfection exclusion

**DOI:** 10.1101/2025.01.26.634807

**Authors:** Anna Sims, Daniel J. Weir, Sarah J. Cole, Edward Hutchinson

**Affiliations:** MRC-University of Glasgow Centre for Virus Research (CVR), Glasgow, UK

## Abstract

The coinfection of individual cells is a requirement for exchange between two or more virus genomes, which is a major mechanism driving virus evolution. Coinfection is restricted by a mechanism known as superinfection exclusion (SIE), which prohibits the infection of a previously infected cell by a related virus after a period of time. SIE regulates coinfection for many different viruses, but its relevance to the infection of Severe Acute Respiratory Syndrome Coronavirus 2 (SARS-CoV-2) was unknown. In this study, we investigated this using a pair of SARS-CoV-2 variant viruses encoding distinct fluorescent reporter proteins. We show for the first time that SARS-CoV-2 coinfection of individual cells is limited temporally by SIE. We defined the kinetics of the onset of SIE for SARS-CoV-2 in this system, showing that the potential for coinfection starts to diminish within the first hour of primary infection, and then falls exponentially as the time between the two infection events is increased. We then asked how these kinetics would affect the potential for coinfection with viruses during a spreading infection. We used plaque assays to model the localised spread of SARS-CoV-2 observed in infected tissue, and showed that the kinetics of SIE restrict coinfection, and therefore sites of possible genetic exchange, to a small interface of infected cells between spreading viral infections. This indicates that SIE, by reducing the likelihood of coinfection of cells, likely reduces the opportunities for genetic exchange between different strains of SARS-CoV-2 and therefore is an underappreciated factor in shaping SARS-CoV-2 evolution.

## Introduction

In 2019, the World Health Organization (WHO) was alerted to a number of cases of fatal viral pneumonia in Wuhan, China [1,2]. This disease, which came to be known as coronavirus disease 19 (COVID-19), was subsequently shown to be caused by severe acute respiratory syndrome coronavirus 2 (SARS-CoV-2). The WHO rapidly declared the disease as a worldwide public health emergency and, as of July 2024, there is approaching 800 million confirmed cases of COVID-19, and over 7 million deaths resulting from the disease [3]. Since its emergence SARS-CoV-2 has continued to circulate and acquire mutations, leading to the generation of numerous variants of concern (VOCs). There is evidence that some VOCs, such as XBB (an omicron derived variant which arose in late 2022), emerged due to recombination, or the exchange of genetic information between different circulating strains [4]. Furthermore, recombinant hotspot analysis has revealed breakpoints around the major coronavirus glycoprotein spike (S) gene, implying that SARS-CoV-2 viruses can effectively to swap Spike (S) genes via recombination [5]. As most neutralising antibodies target the S protein following natural infection and vaccination [6], swapping S genes may increase the ability of SARS-CoV-2 to spread in an otherwise immune population. Therefore, recombination between strains of SARS-CoV-2 and the factors controlling the likelihood of recombination, remains a concern for public health [7].

Recombination between viruses requires the coinfection of cells [8,9]. This is regulated, in part, by superinfection exclusion (SIE), which blocks secondary infection of a cell which had been previously infected with a related virus after a period of time. SIE has been demonstrated to be active during the infection of cells with many viruses of bacteria, plants and animals [10–23], which is achieved through a variety of different mechanisms [24–28]. In a recent study, Bonavita *et al.* provided evidence for active SIE mechanisms between two genetically distinct coronaviruses, canine coronavirus (CaCoV) and feline coronavirus (FeCoV) [8]. The group showed that infecting cells with CaCoV 2 hours before infection with FeCoV, reduced the replication of FeCoV 20-fold. Increasing the time between infection events to 4, 8 and 12 hours (h) only increased the effect on FeCoV replication. Interestingly the group found no suppression of CaCoV when FeCoV was used as a primary infecting virus, until the time between infection events was increased to 12 h. The difference in timing of onset of SIE hints to different mechanisms being employed by FeCoV and CaCoV. [24–28]. In this study, we explored the possibility of active SIE mechanisms during SARS-CoV-2 coinfection, to our knowledge the first time this has been investigated for this clinically important coronavirus.

Aside from SIE, the frequency of cellular coinfection and therefore recombination can also be controlled by the localised spread of infections [29]. This is because if viruses within an organ are confined to spread locally to adjacent cells, the likelihood of viruses from different areas coinfecting the same cell are low. We have shown previously that SIE restricts coinfection to small interface regions during localised spread of influenza viruses, thereby likely limiting the opportunity of genetic exchange between viruses [30]. We hypothesise that active SIE mechanisms would similarly affect SARS-CoV-2 during localised viral spread.

There is evidence of localised viral spread within the organs of SARS-CoV-2 infected patients and experimentally infected animals. Antigen in post-mortem samples cluster to form foci in tissues of the lung, small intestine, and kidney [31], though clustering can be partially lost later in the course of disease [32–34]. Experimental infection of human lung and bronchi tissues *ex vivo*, and human epithelial cell cultures show distinct SARS-CoV-2 N protein-positive foci with multiple SARS-CoV-2 variants (Alpha, Beta, Delta and Omicron) [35–37]. Foci have also been observed in the respiratory epithelium of rhesus macaques and ferrets following experimental infections [38,39]. One study utilised light sheet microscopy to visualise SARS-CoV-2 foci in nasal turbinates of ferrets at day 4 post infection [40]. Using this technique, they could not only image individual foci in whole lungs but measure the linear distance between the foci in micrometres (between 555-1307 µm), revealing the distance the viruses must spread before they are able to coinfect cells. Therefore, all the evidence we have of direct visualisation of infection in lungs show discrete foci formed by localised SARS-CoV-2 spread, but how this impacts the likelihood of coinfection has not been investigated. We therefore hypothesised that, as SARS-CoV-2 forms localised infected regions, if SIE regulates SARS-CoV-2 coinfections at a cellular level it would also, as for influenza A viruses, limit the opportunities for SARS-CoV-2 coinfection in a spreading infection.

Here, using isogenic SARS-CoV-2 viruses encoding fluorescent reporters we describe, to our knowledge for the first time, that SIE restricts SARS-CoV-2 coinfection of cells. We show that the potential for coinfection of cells is reduced within the first hour of SARS-CoV-2 infection, and then reduces exponentially as the time between primary and secondary infection is increased. Applying this to the localised spread of SARS-CoV-2 infections by modelling foci in 2D using plaque assays, we show that SIE restricts coinfection of SARS-CoV-2 spreading from different regions of infection, limiting the likelihood for genetic exchange between viruses. Our results show that SIE is an underappreciated barrier to SARS-CoV-2 coinfection, and we conclude that by restricting opportunities for genetic exchange, SIE is likely to impact the ongoing evolution of SARS-CoV-2 viruses.

## Results

### Quantifying coinfection of cells with SARS-CoV-2 using microscopy and flow cytometry

It has been reported from multiple groups that the SIE acts between closely related coinfecting viruses, and the effect is lost when the viruses are unrelated to each other [14,16,18,23,41]. Therefore, in order to maximise the chance to observe SIE induced by SARS-CoV-2, we chose to investigate this using two isogenic variants of SARS-CoV-2 reporter viruses, which differ only in the fluorophore they encode. To do this we used an established SARS-CoV-2 reporter virus system [42]. This system consists of isogenic SARS-CoV-2 viruses based on the Wuhan-Hu-1 isolate which encode a fluorescent protein (in this study; ZsGreen or mCherry) downstream of the *ORF7a* gene (which encodes the NS7A protein) [43,44].

Aside from viral relatedness, the replication rate of the primary infecting virus has also been hypothesised to influence SIE kinetics, with a faster the replication rate resulting in quicker onset of SIE [8,30]. As the reporter viruses we used were isogenic, aside from the gene encoding the fluorescent protein, we expected that the viruses would replicate at similar rates to each other in VAT cells (ACE2 and TMPRSS2-overexpressing VeroE6 cells), which have been shown to be susceptible to SARS-CoV-2 infection [42]. Using classical plaque assays, we found that the viruses replicated with similar kinetics (fig. 1). Despite a lower mean titre for SARS-CoV-2 mCherry than SARS-CoV-2 ZsGreen at 24 hpi, there was no significant difference between the titres of each virus at any timepoint (paired Wilcoxon non-parametric test, p>0.05).

**Figure 1:**
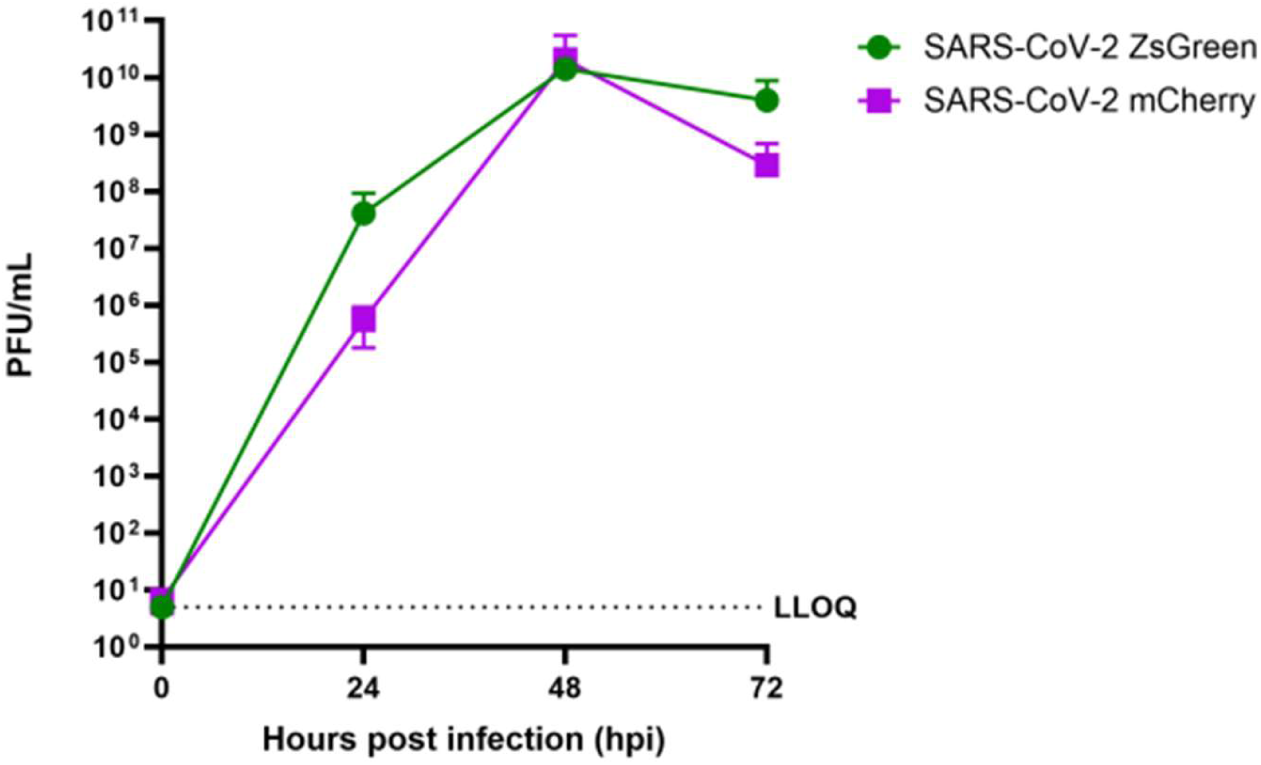
SARS-CoV-2 reporter viruses replicate with similar kinetics. Multicycle growth kinetics of SARS-CoV-2 reporter viruses were assessed by infecting VAT cell monolayers with viruses at a multiplicity of infection (MOI) of 0.01 plaque forming units per cell (PFU/cell), and harvesting the supernatant at the timepoints indicated. Virus titres were then calculated from plaque assays on VAT cells. The data represent the mean and standard deviation of 3 independent repeats. The lower limit of quantification (LLOQ) is represented by a dashed line. Differences between SARS-CoV-2 ZsGreen and mCherry growth were non-significant (P>0.05) at each timepoint as assessed using a paired, non-parametric Wilcoxon test.

Following this, to confirm the expression of the reporter gene in VAT cells, we infected the cells on coverslips with 0.5 PFU/cell of each virus and imaged at 24 hours post infection (hpi) using a high-resolution confocal microscope. We observed singly infected cells (positive for either magenta, indicating infection with SARS-CoV-2 mCherry) or green fluorescence, indicating infection with SARS-CoV-2 ZsGreen), and uninfected cells (where only the nuclear DAPI stain can be seen); (fig.2A). Similarly, we showed that we can separate ZsGreen and mCherry expressing cells in flow cytometric analysis at 16 hpi (fig.2B). Therefore, the SARS-CoV-2 reporter viruses are suitable for our investigation of SARS-CoV-2 SIE.

**Figure 2:**
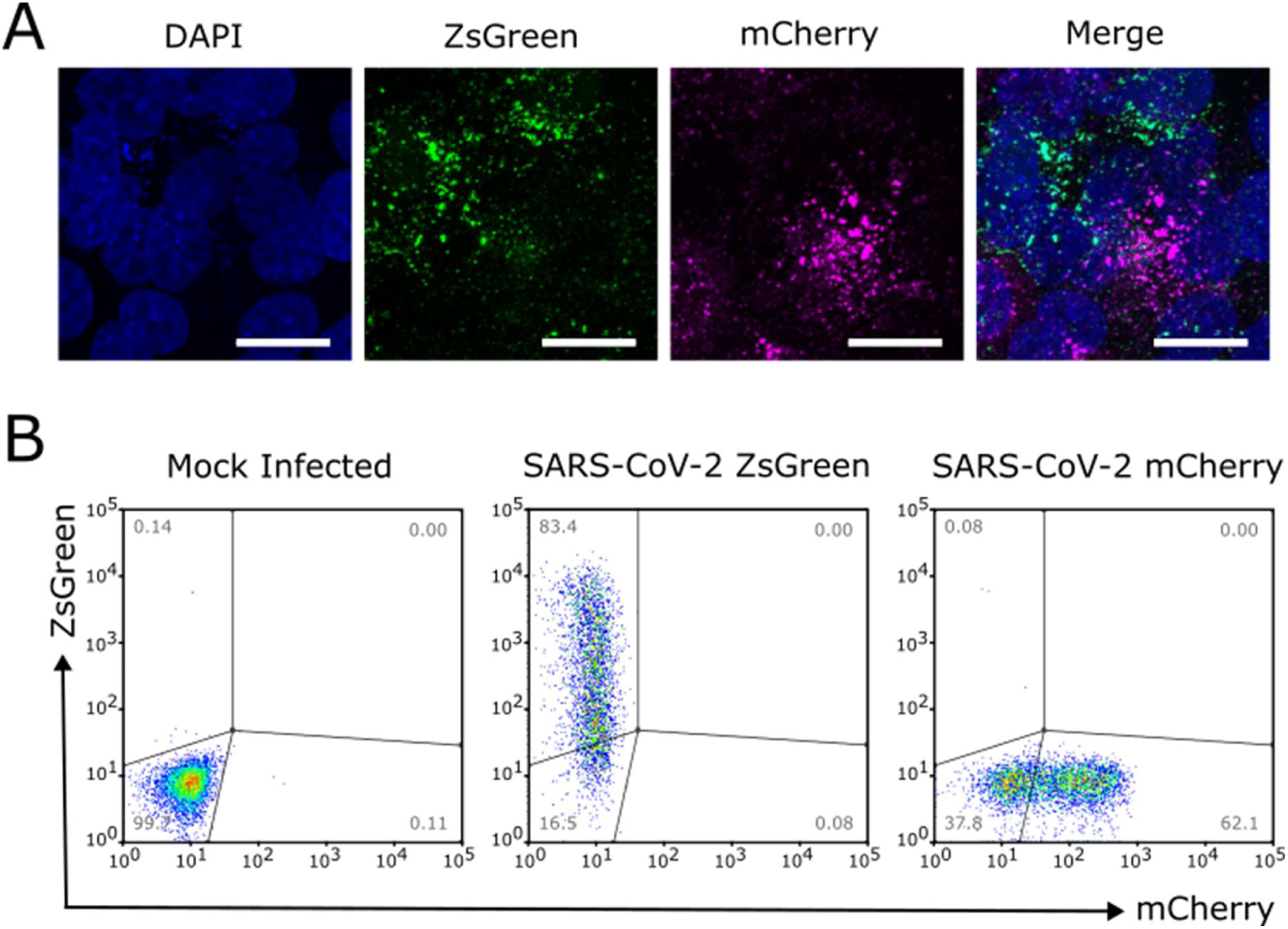
Cells infected with SARS-CoV-2 reporter viruses can be distinguished by confocal microscopy and flow cytometry. (A) Images of cells infected with SARS-CoV-2 reporter viruses taken using a high-resolution confocal microscope. VAT cells were simultaneously infected at an MOI of 0.5 PFU/cell of each virus, and fixed at 24 hpi. Images were obtained using a 63× objective. Scale bar = 20 μm. (B) Flow cytometric analysis of VAT cells separately infected with SARS-CoV-2 reporter viruses. Cells were infected at an MOI of 2 fluorescence forming units per cell (FFU/cell) of either ZsGreen or mCherry expressing viruses, and harvested for analysis 16 hpi. Gate frequencies as percentage of total cells are shown.

### SIE inhibits coinfection of cells with SARS-CoV-2

It was not known whether SARS-CoV-2 infection causes SIE onset in cells. We investigated this in using our fluorescent reporter virus system in VAT cells. To do this we first infected monolayers of cells at various time intervals with SARS-CoV-2 ZsGreen and then subsequently infected them with SARS-CoV-2 mCherry. The experimental procedure is graphically represented in fig.3A. We then harvested the cells at 16 hpi and measured the expression of the fluorophores using flow cytometry. We used an MOI of 2 FFU/cell for both viruses in order to ensure that nearly all cells would be infected, and as expected, infection with this amount resulted in the majority of cells expressing the corresponding fluorescence (96.4% and 87.5% cells positive for ZsGreen and mCherry respectively) during infection with the viruses alone (fig.3B).

**Figure 3:**
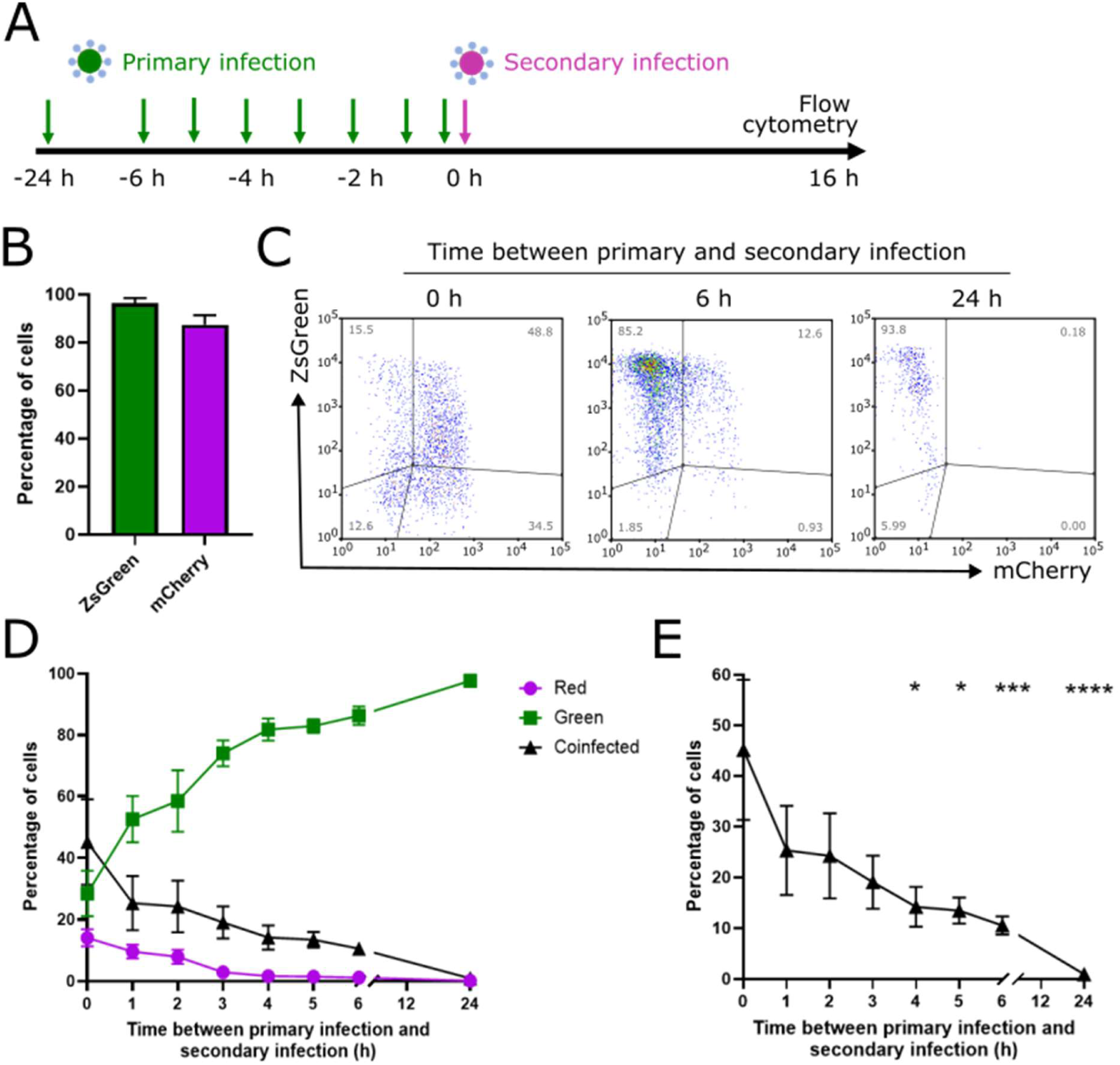
SIE restricts infection of SARS-CoV-2 mCherry in VAT cells previously infected with SARS-CoV-2 ZsGreen. (A) Schematic of the experimental investigation of SARS-CoV-2 SIE. (B) Percentage of positive cells during individual infection of SARS-CoV-2 ZsGreen and mCherry reporter viruses. VAT cells were infected with MOI 2 FFU/cell at the time points indicated and harvested for flow cytometry at 16 hpi. Data represents mean and SD, n=2. (C) Representative flow cytometry plots of cells infected with reporter viruses. VAT cells were first infected with SARS-CoV-2 ZsGreen, before secondary infection at the time points indicated with SARS-CoV-2 mCherry. (D) Kinetics of onset of SIE determined by flow cytometric analysis. Data represents mean and SD, n=5. (E) Percentage of coinfected cells as the time between infection events is increased. Significance determined by non-parametric, Friedman multiple comparison test, *p<0.05, ***p<0.0005, p<0.0001.

In coinfection scenarios, we gated the cells into different populations based on the expression of both fluorophores (fig.3C). We found that the proportion of coinfected cells reduced steadily as the time between primary and secondary infection was increased (fig.3D), which became statistically significant by 4 hours between infection events (p=0.0047, Friedman multiple comparisons test), and was significant thereafter (fig.3E). The cells remain resistant to secondary infection at 24 h between infection events, the longest delay tested, by which point the proportion of coinfected cells had reduced to nearly 0. It is worthy of noting however, that the number of cells collected by the flow cytometer at 24 h was lower than the other conditions tested (fig.3C). This is likely due to the cells dying and therefore being lost from the analysis, but it demonstrates the cells which remain alive after 24 h remain resistant to secondary infection. Interestingly, although the viruses used in this study are isogenic, when we performed the reciprocal experiment (primary infection with SARS-CoV-2 mCherry and secondary infection with SARS-CoV-2 ZsGreen), we found subtle differences in the kinetics of SIE onset (Figure 4). Generally, both experiments follow the same trend (a reduction in the amount of coinfected cells) but with SARS-CoV-2 mCherry applied first, the reduction does not become significant until there was 6 h gap between primary and secondary infection (p=0.0044, Friedman multiple comparisons test). We hypothesise that these differences may be due to (non-statistically significant) differences in the early replication rate (before 24 hpi) between SARS-CoV-2 ZsGreen and SARS-CoV-2 mCherry (fig. 1).

**Figure 4:**
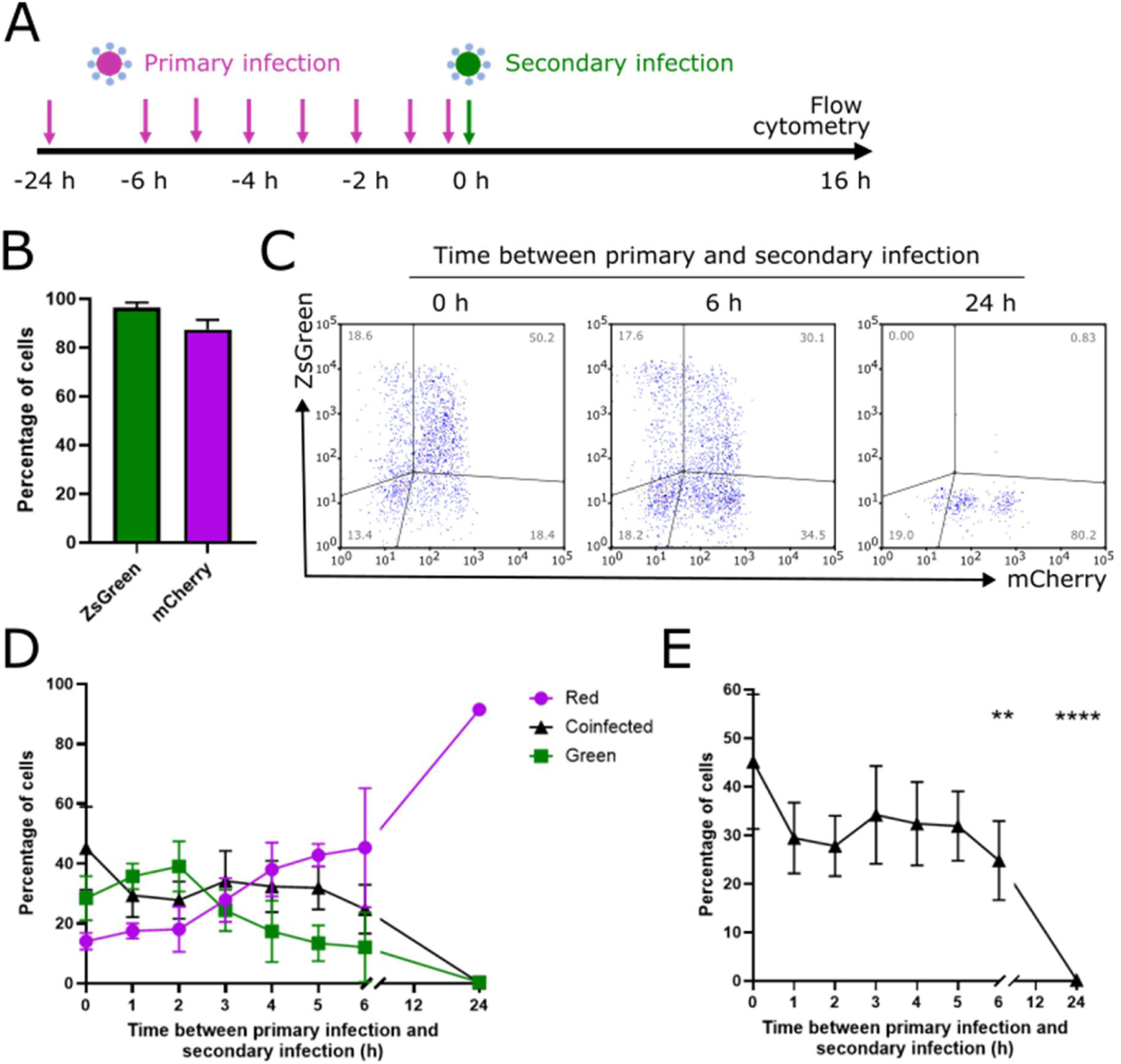
SIE restricts infection of SARS-CoV-2 ZsGreen in VAT cells previously infected with SARS-CoV-2 mCherry. (A) Schematic of the experimental investigation of SARS-CoV-2 SIE. (B) Percentage of positive cells during individual infection of SARS-CoV-2 ZsGreen and mCherry viruses. Measurement of singly infected VAT cells with MOI 2 FFU/cell tagged viruses harvested for flow cytometry at 16 hpi. Data represents mean and SD, n=2. (C) Representative flow cytometry plots of cells infected with reporter viruses. VAT cells were first infected with SARS-CoV-2 mCherry, before secondary infection at the time points indicated with SARS-CoV-2 ZsGreen. (D) Kinetics of onset of SIE determined by flow cytometric analysis. Data represents mean and SD, n=5. (E) Percentage of coinfected cells as the time between infection events is increased. Significance determined by non-parametric, Friedman multiple comparison test, *p<0.05, ***p<0.0005, p<0.0001.

Overall, we do observe a reciprocal onset of SIE between SARS-CoV-2 ZsGreen and mCherry in VAT cells. Therefore, our data offers the first description of SIE initiated by SARS-CoV-2 infection and displays a time dependent resistance to coinfection.

### SIE onset exponentially reduces in the ability of the secondary virus to infect previously infected cells

We wished to more closely examine the kinetics of the onset of SIE in our system, as this may give us insights into the mechanism(s) of SIE for SARS-CoV-2. To do this, we used the proportion of cells that expressed the reporter of the secondary infecting virus (mCherry in fig. 5A, and ZsGreen in fig. 5B) (including both singly and coinfected cells), and from this calculated the titre of the productively infecting viruses to give red forming units (RFU) and green forming units (GFU) per ml of virus. To do this we assumed that viruses applied at the same time had equal opportunity to infect the cells, and therefore the infection could be modelled by a Poisson distribution (see materials and methods for details). We note that, as the entry of the primary infecting virus was not synchronised in the previous experiments, each individual cell is likely to have different kinetics of SIE onset. Our analysis describes the average SIE onset across a group of cells rather than the kinetics of SIE onset with any one single cell. As expected, we observed a decrease in the ability of the secondary infecting virus to productively infect the previously infected cells, which became more pronounced as the time between the infection events increased (fig.5).

**Figure 5:**
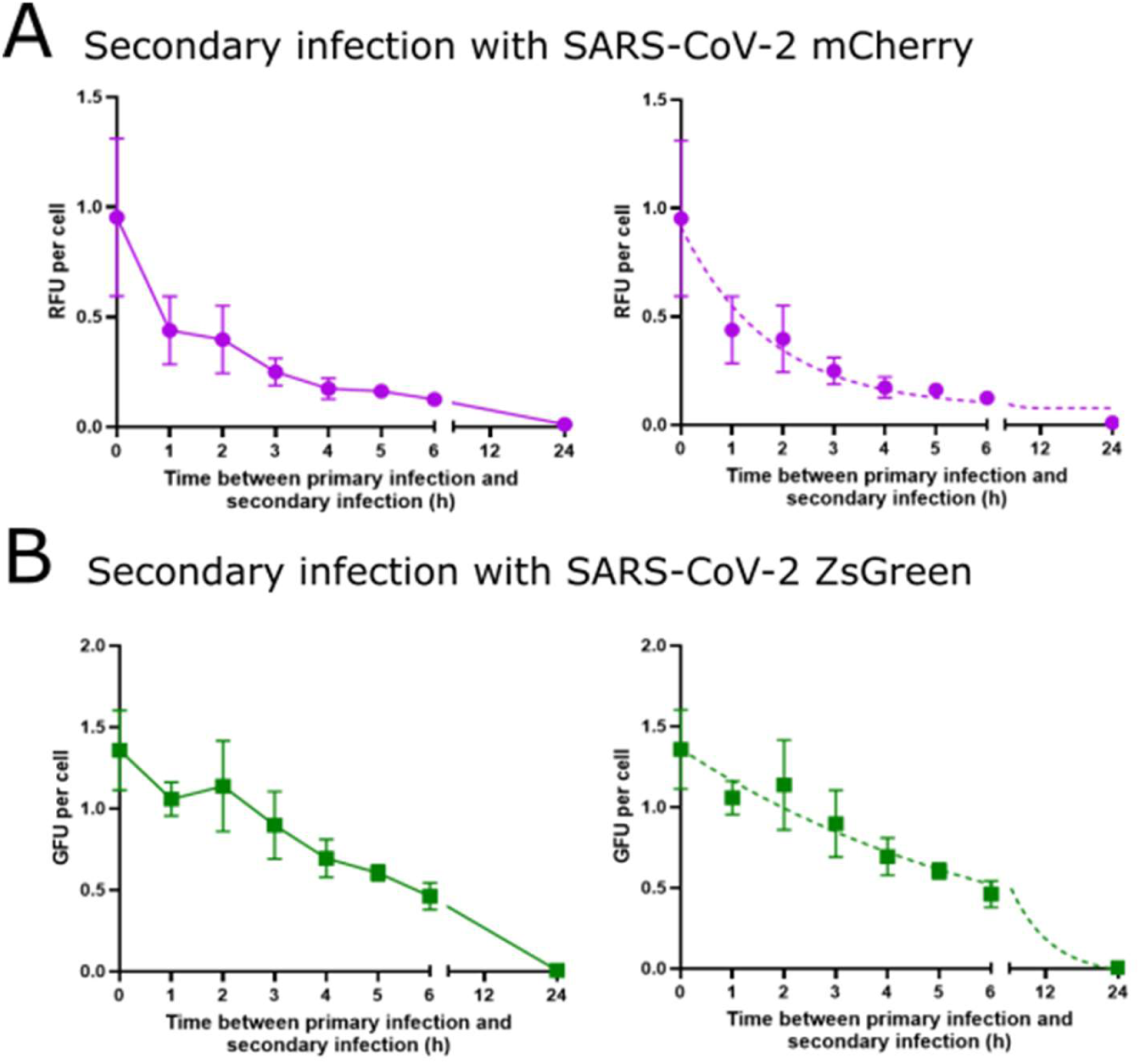
Kinetics of onset of SIE by SARS-CoV-2 can be described by an exponential decay model. The kinetics of SIE onset were calculated from data in figures 3 and 4 when the secondary virus was (A) SARS-CoV-2 mCherry or (B) SARS-CoV-2 ZsGreen. RFU and GFU per cell were calculated from the proportions of green, red and coinfected cells (see text for details). Data points represents the mean and SD (n=5). Solid line represents line of best fit, dotted line represents exponential decay model fit.

We hypothesised that SIE onset in SARS-CoV-2 infected cells might be linked to the replication of the primary infecting virus. If this was the case, we would expect that the kinetics of SIE onset would occur exponentially, following the kinetics previously reported for the synthesis of SARS-CoV-2 RNA following infection [45,46]. To investigate this, we applied a one phase decay model to the RFU/cell (fig. 5A) and GFU/cell (fig. 5B), with the plateau constrained above 0. The model was fitted by least squares. We found the model fit the data with reasonable fitness (total sum of squares (SST) = 0.85 and 1.05 respectively). As previously mentioned, the kinetics of SIE onset differed depending on the primary infecting virus used, which is reflected in the half-life of the model (1.2 h for SARS-CoV-2 ZsGreen and 4.5 h for SARS-CoV-2 mCherry). Although the underlying mechanism of SIE in SARS-CoV-2 remains to be determined, the consistency between our observations and an exponential decay model suggests that it is likely to be directly or indirectly connected to transcription or replication of the initial SARS-CoV-2 genome to infect the cell.

### SIE inhibits coinfection between SARS-CoV-2 viruses during localised viral spread

Having determined that SIE restricts coinfection between sequentially delivered variants of SARS-CoV-2 at the single cell level, we set out to investigate whether it restricts coinfection between viral populations during localised viral spread across a monolayer of cells. To do this, we performed a classical plaque assay, growing VAT cells to near-confluency before infection, and limiting long-range viral spread using a semi-solid overlay. This was done in order to recapitulate the highly spatialised spread of SARS-CoV-2 that has been observed within SARS-CoV-2 infected patients and experimentally infected animals [36,37,40]. We decided to investigate the potential for coinfection between viruses located in different foci of infection, as this is the mostly likely scenario which could lead to recombination between different lineages of SARS-CoV-2, and therefore the most interesting scenario for recombination. To do this, we infected VAT monolayers with a mixture of ZsGreen and mCherry viruses and allowed foci of infection to form under a 0.6% (w/v) Avicel overlay in viral growth media (VGM). At 24 hpi we removed the Avicel, fixed the cells and imaged using fluorescent microscopy.

We observed that (as is typical under plaque assay conditions) infected foci grow with roughly circular profiles (fig.6A). However, where foci infected by of SARS-CoV-2 mCherry and ZsGreen meet, these circular profiles are sharply interrupted with minimal overlap. The profile of these interacting foci strongly suggests that the presence of each infected area blocked the other from expanding (fig.6B). In order to more closely investigate this overlap we used FIJI ImageJ [47] to apply a binary threshold to the fluorophore expression, and calculated the percentage of pixels in each image which displayed red and green together, or separately, above this threshold. We observed only small areas of co-expression (denoting coinfected cells) at the boundaries where red and green foci met (fig.6C). On average, this coinfected region constituted only around 2.9% of the infected area in each image (fig.6D). This suggests that SIE limits the potential for coinfection between SARS-CoV-2 populations in different foci of infection, the scenario most likely to lead to recombination between distinct lineages of SARS-CoV-2 within the respiratory tract.

**Figure 6:**
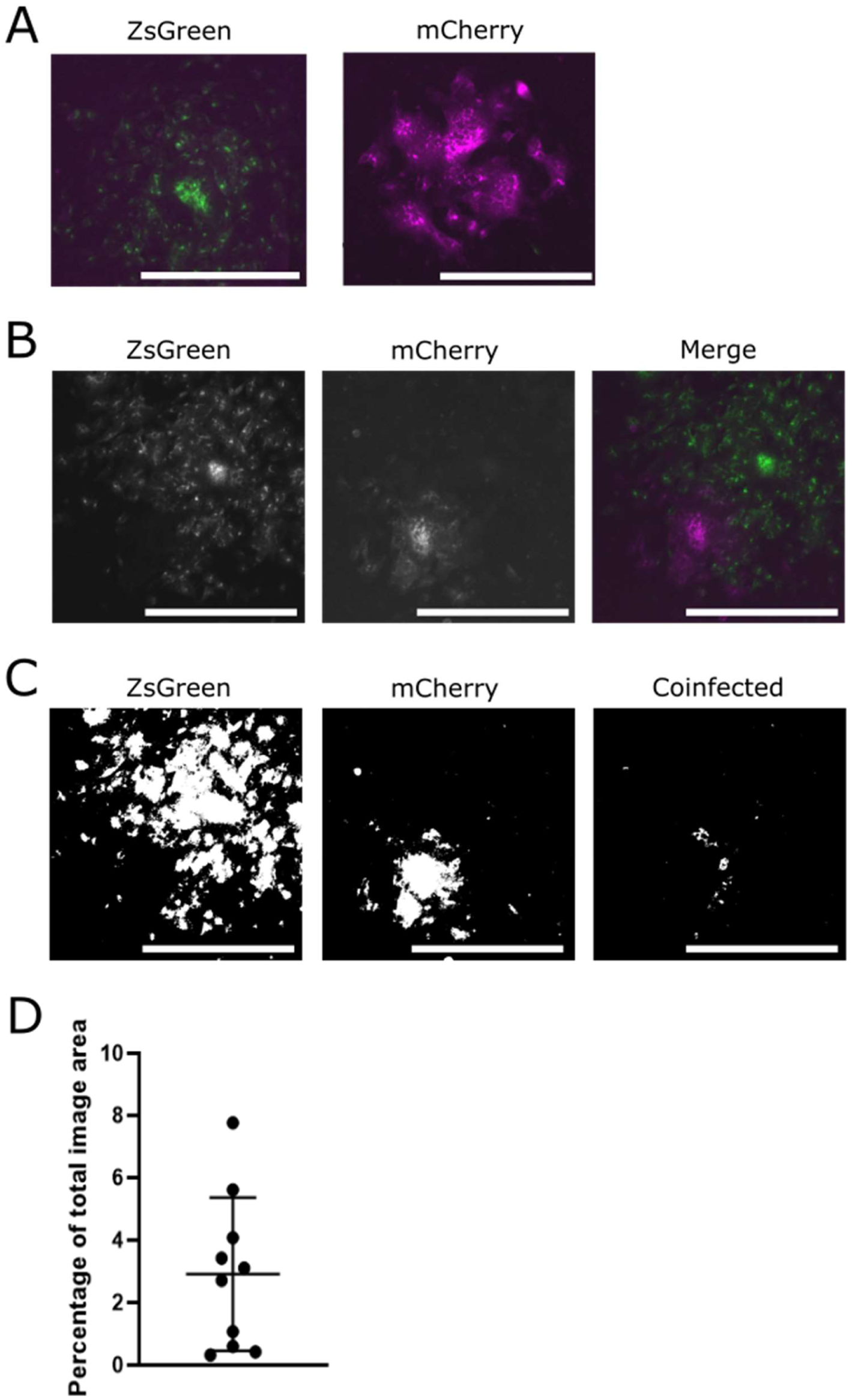
SIE spatially restricts interactions between viruses from separate SARS-CoV-2 foci of infection. Monolayers of VAT cells were infected with ZsGreen and mCherry tagged SARS-CoV-2 viruses, overlayed with 0.6% Avicell, fixed at 24 hpi and imaged using Celigo fluorescent microscope. Scale bar = 250 µm. (A) Representative images of plaques, merged images of green and red channel (B) Representative image of foci interaction, single channel images and merged channel image (C) A binary threshold was applied to images of foci to distinguish cells expressing either the ZsGreen, mCherry, or both fluorophores together. Threshold applied to the image in (B) is shown. (D) The percentage of coinfected areas in comparison to total image area was calculated from images at taken at 24 hpi. The mean and SD of 10 individual fields of view from 1 experiment are shown.

## Discussion

In this study, we provide the first experimental evidence that coinfection of cells with SARS-CoV-2 is limited by SIE. We also show that the exponential onset of SIE restricts the opportunity for coinfection, and therefore likely genetic exchange, between SARS-CoV-2 viruses during a model of localised viral spread. Our data indicate the SIE is likely to control the potential for recombination events between distinct lineages of SARS-CoV-2, which is an ongoing concern for public health.

The kinetics of SIE onset in SARS-CoV-2 infections were a good fit for an exponential model, which in turn fits with the accumulation of newly synthesised viral RNA in the cell [45,46]. Although both viruses induced SIE in an exponential fashion, we observed minor variations in the parameters of this model even with these isogenic viruses, which suggests that the parameters of the exponential onset of SIE may well vary for strains and cell types where SARS-CoV-2 replication proceeds at different rates. As we measured SIE using the expression of a reporter protein encoded by the secondary infecting virus, we can conclude that for SARS-CoV-2 SIE affects a step in viral replication before the translation of viral proteins. While the underlying mechanism for SIE in coronaviruses is not yet known, this correlation between SIE onset and the replication rate of the primary infecting virus has been observed in other coronaviruses, namely canine coronavirus (CaCoV) and feline coronavirus (FeCoV) [8] as well as in unrelated viruses, for example influenza

A viruses [12], and suggests that the underlying mechanism for SIE may be directly or indirectly connected to viral replication. The link between the length of the virus lifecycle and the kinetics of SIE onset has not been fully explored. For mechanisms where a particular viral protein is required for SIE onset (such as human immunodeficiency virus 1 (HIV-1) Nef protein binding to and inducing the internalisation of the viral receptor molecule, CD4 [13,24]), it is clear how a faster viral replication cycle would induce faster onset of SIE. For SARS-CoV-2, the mechanism of SIE and its relation to the progress of the viral replication cycle remains to be determined

An alternative explanation for the onset of SIE would be the innate immune response to infection, however the cell line we used (VAT cells) do not secrete interferon alpha or beta (IFN) [48], indicating that the SIE observed here did not dependent on a type 1 IFN response. SIE in Vero cells has also been demonstrated following infection with influenza A viruses [12] and rat borna viruses [49]. Our finding therefore adds to the increasing evidence that SIE is not driven primarily by innate immune mechanisms following viral infection [23,41]. It is important to note that the cellular immune system is highly redundant, and therefore other innate immune mechanisms may be acting to compensate for the blocked pathway, both in this study and in others [50]. Of note, is that despite being deficient in type 1 IFN response, Vero cells have been shown to secrete type 3 IFNs during infection, which are capable of upregulating key interferon stimulated genes (ISGs) and inducing a refractory state in recipient cells [51]. It is also possible that the kinetics of SIE would be altered if the type 1 interferon response was active, and that innate and non-innate mechanisms could work together to induce SIE. Further work will be needed to systematically assess how different aspects of innate immunity could modulate the rate of SIE onset in SARS-CoV-2.

Having identified SIE for SARS-CoV-2 and defined its kinetics at a cellular level, we asked how it could affect coinfection between spreading virus populations, the most likely scenario for genetic exchange between SARS-CoV-2 strains within a host. This is a reasonable scenario for a natural coinfection, as even if the strains were acquired by a host organism at a similar time, the chance that the viruses reach the same cell at the same time are vanishingly small, considering the number of cells in the respiratory tract, and the small size of the transmission bottlenecks for SARS-CoV-2 [52,53]. We therefore expect recombination to occur after the viruses establish an infection and meet another focus of infection while spreading through the respiratory tract. We showed in this study that coinfection in this scenario will be severely constrained by SIE. In a previous study, we showed the same effect occurs during localised spread of influenza A viruses both *in vitro* and *in vivo* [30]. We propose that constraint by SIE is a common feature of the interactions between respiratory viruses as they spread locally within the respiratory epithelium.

Together our results highlight that SIE is an underappreciated barrier to coinfection and therefore recombination of SARS-CoV-2. Recombination is an important aspect of coronavirus biology. There have been suggestions that historical recombination between related coronaviruses may have led to the emergence of the original SARS-CoV-1 in 2003 [54]. Similarly, there is evidence for recombination in the evolutionary history of SARS-CoV-2 [55]. During the COVID-19 pandemic, recombinant lineages of SARS-CoV-2 were found to be circulating in the United Kingdom in late 2020 and early 2021 [56]. Recombination was even detected within a single individual superinfected with different variants (Alpha and Epsilon) of SARS-CoV-2, with recombination detected in S, N and ORF8 coding regions of the genome [57]. However, there is evidence that when the global population of viruses is considered recombination in SARS-CoV-2 is quite rare. Sequencing studies noted a lack of recombinant variants circulating during the pandemic [58,59]. Recently, Bonavita et al described recombination between coronaviruses as “both frequent and rare”, in that if coinfection occurs in an experimental setting, recombinant RNAs were always detected, but that they represent a surprisingly low proportion of the RNA species produced [8]. Most of the experiments were performed with viruses administered simultaneously, but when infection events were staggered, the group observed SIE onset, and subsequently observed a reduction in the detection of recombinant RNA species. This provides powerful evidence that coinfection potential, as regulated by SIE, is the primary gatekeeper of recombination between coronaviruses. We anticipate, based on these results and ours, that SIE represents a meaningful barrier to the generation of recombinant VOCs. Whether we can harness this mechanism to prevent emergence of recombinant VOCs is currently unexplored.

Overall, this study confirms for the first time that SIE, a common feature of infection by viruses from many diverse viral families, is active in the infection of cells by SARS-CoV-2. We explore how this shapes the interaction dynamics between SARS-CoV-2 viruses during localised viral spread including reducing the likelihood of recombination between viruses. More research is required if we are to meaningfully understand the role of SIE in shaping the ongoing adaption of SARS-CoV-2, and the mechanisms underlying this process.

## Materials and Methods

### Cells and viruses

VeroE6-ACE2-TMPRSS2 (VAT) cells (a gift from Prof. S Wilson, University of Cambridge) were maintained in complete media (Dulbecco’s Modified Eagle Medium (DMEM, Gibco) supplemented with 10% Foetal Bovine Serum (FBS, Gibco)). All cells were maintained at 37 °C and 5% CO^2^ in a humidified incubator.

SARS-CoV-2 reporter viruses (a gift from Dr. S. Rihn, University of Cambridge) were rescued as previously described [42]. The viruses were passaged at low MOI in viral growth media (VGM) (DMEM with 2% v/v FBS) to create a working stock. Viral variants were propagated separately and were only used together in coinfection experiments indicated in the figure legend. All handling of SARS-CoV-2 viruses was performed in a biosafety containment level 3 (CL3) laboratory.

Viral infectious titre in plaque forming units per millilitre (PFU/mL) was obtained from VAT cells using standard conditions. Briefly, viruses were serially diluted in VGM and used to infect confluent cell monolayers for 1 hr at 37°C before overlay with 0.6% (w/v) avicell in VGM. The plates were incubated for 72 h at 37°C before fixation and staining with Coomassie Blue staining solution (0.1% Coomassie Brilliant Blue R-250/45% methanol/10% acetic acid), and the plaques calculated.fo

In some case as indicated by the figure legend, virus infectious titre in fluorescence forming units per millilitre (FFU/mL) was used. Viruses were serially diluted in VGM, and inoculated onto confluent VAT cell monolayers for 1 h at 37°C before overlay with complete medium. At 24 h the proportion fluorescence positive cells by flow cytometry (see below) and the FFU/cell calculated using the following formulae:

When the proportion of positive cells (P) was measured and P(0<X>1), to calculate the MOI in

FFU/cell *m:*

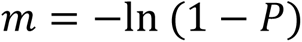

Thus, to calculate the virus titre (FFU/mL):

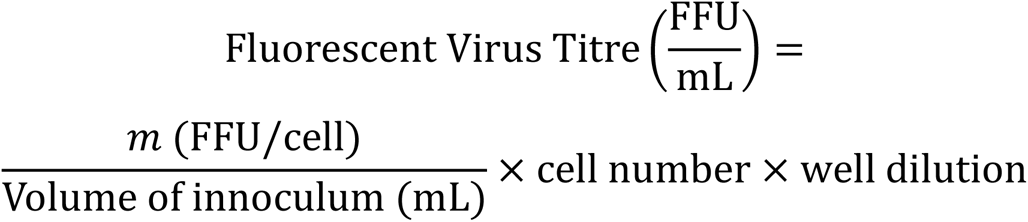

### Immunofluorescence and Imaging

Confocal images of SARS-CoV-2 infected cells were obtained by infecting confluent VAT cells on coverslips, with an MOI of 0.5 PFU/cell for each of the SARS-CoV-2 viruses, for 24 hours before fixation in 8% (v/v) formaldehyde diluted in PBS (Sigma-Aldrich).

Following fixation, the cells were rinsed in PBS and the nucleus stained with 4′,6-diamidino-2-phenylindole (DAPI). Coverslips were then mounted and imaged. Confocal microscopy was performed using the Zeiss LSM 880 (63x oil immersion objective, 1.4 NA). Super resolution imaging of coinfected cells was performed by performing Z-stacks encompassing 3D regions of interest using Airyscan fast detection. Post-acquisition auto airyscan processing was performed within Zen Black (Zeiss) software (v14.0.29.201). Maximum intensity projections of 3D airyscan processed Z-stacked images were created using the Zen Black (Zeiss) software (v14.0.29.201). Mock, or single infected cells were imaged similarly but in 2D. Image intensity values were adjusted according to mock infected controls.

### Foci interaction assay

To obtain images of interactions between SARS-CoV-2 foci of infection, VAT cell monolayers were infected with a diluted mixture of mCherry and ZsGreen tagged viruses after which a 0.6% (w/v) avicell overlay in VGM was applied and incubated at 37°C for 24 h before fixation for in 8% (v/v) formaldehyde for 1 h. Following this the cells were rinsed in PBS and imaged using the Celigo imaging cytometer (Nexcelom). Images were processed in FIJII ImageJ [47] using custom macros which can be accessed here: https://github.com/annasimssarscov2/SARSSIE

### Viral Growth Kinetics

For SARS-CoV-2 multicycle growth kinetics, viruses were applied to confluent VAT monolayers at an MOI of 0.001 PFU/cell and the cells were incubated with the inoculum for 1 h at 37°C to allow the viruses to enter cells. Following this, the inoculum was removed, and fresh VGM was added. Media were sampled at the time points indicated, and stored at −80°C before titration by plaque assay.

### Flow cytometry

At various time intervals, VAT cells were inoculated with reporter viruses, diluted in VGM at MOI of 2 FFU/cell for 1 h at 37°C. After this the inoculum was removed and replaced with complete media. After this, cells were inoculated for 1 h with a second reporter virus, also at MOI of 2 FFU/cell. After 1 h this inoculum was removed and replaced with complete media, and the cells were incubated for a further 16 h at 37°C.

The proportions of cells expressing the different fluorophores were assessed using a Guava easyCyte HT System cytometer (Luminex). Briefly, infected and mock-infected cell monolayers were dissociated in TrypLE express (Gibco) for 15 minutes at 37°C and dispersed into a single-cell suspension before fixation in 8% formaldehyde (v/v) in PBS.

Flow cytometry data were analysed in FlowJo software v10.6. The thresholds for assessing positive detection of the red and green fluorophores were set using the mock-infected cells as a negative control.

### Analysis and Modelling

Statistical analysis was carried out using Graphpad Prism (version 9.1.0). Statistical tests where applicable are described in the figure legends. Data were visualised using Graphpad Prism version 9.1.0 or FIJI ImageJ [47].

We calculated the titre of viruses that would cause that proportion of cells to become fluorescent (either green fluorescent units, GFU or red fluorescent units RFU) using the Poisson distribution [60]. By using the Poisson distribution, we make the assumptions that (i) viruses that are added to cells at the same time will infect independently of each other; and (ii) at the point that the primary virus is added, all cells are equally permissive to infection. Therefore, by rearranging the Poisson distribution formula we calculated RFU and GFU as follows:

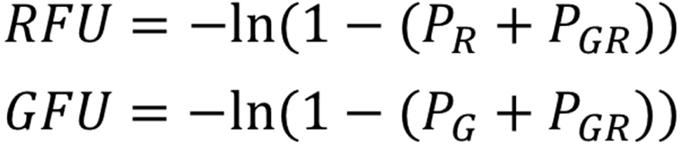

Whereby *P*_R_ is the proportion of cells that express only the red fluorophore, *P*_G_ is the proportion of cells expressing only the green fluorophore and *P*_GR_is the proportion of cells that expresses both fluorophores.

A one phase decay model was fitted to these data by least squares, using GraphPad Prism (version 9.4.1; GraphPad. The plateau of the model was constrained above 0 RFU/cell or GFU/cell.

## Acknowledgements

We would like to thank Dr A. Wickenhagen and Dr M. Turnball for their help and advice in handling the fluorescent SARS-CoV-2 viruses used in this study. Additionally, we would like to thank Dr. A. Szemiel and E. Davis for their help in training A.S. for work at CL3.

## Author Contributions

A.S. investigation, conceptualisation, methodology, visualisation, analysis, writing – original draft; D.W. investigation, visualisation, writing – review and editing; S.C. investigation, writing – review and editing; E.H. supervision, writing – review and editing, funding acquisition.

